# The Gender Gap Amongst Doctoral Students in the Biomedical Sciences

**DOI:** 10.1101/2022.10.18.512765

**Authors:** Michael D. Schaller

**Affiliations:** Department of Biochemistry and Molecular Medicine, West Virginia University School of Medicine, Morgantown, West Virginia 26506

**Keywords:** doctoral students, productivity, gender, publication, citation

## Abstract

Historically women have been underrepresented in STEM careers. While the number of women receiving doctorate degrees in the biological sciences has exceeded the number of men since approximately 2005, there is still a disparity between the sexes at more advanced career stages. Achieving equity is an important social goal and there is an expected benefit to science since diverse groups outperform homogeneous groups. There are many factors that contribute to the disparity between men and women in science, including a disparity in research productivity. While many studies have documented this “productivity paradox” and examined factors driving this disparity, few studies have addressed differences in productivity between men and women doctoral students. This is an important population to assess since the individuals are in the formative stages of their academic career and differences in productivity could have a significant impact on career progression. This study addresses this question and identified more than 42,000 doctoral students in the biological and biomedical sciences working with over 16,000 advisors at 235 institutions in the United States and finds a disparity in research productivity between men and women. Men produce >10% more first author papers, >15% more total papers and their first author papers receive >17% more citations. The findings establish the generality of the gender gap in research productivity among doctoral students in the biological and biomedical sciences. Redressing this gap at the formative stage of young scientists’ careers, when they are establishing their credentials to advance in their field, is important to address the disparity between the sexes in the biomedical workforce.

## Introduction

Despite awareness and efforts to redress diversity in science, women remain underrepresented in academia. The most recent Survey of Earned Doctorates revealed that 53.8% of doctoral recipients in the biological and biomedical sciences in 2020 were women, demonstrating an over representation of women earning PhDs (1). The most recent Survey of Doctoral Recipients revealed that women are underrepresented among the faculty in the biological, agricultural and environmental life sciences at 4 year degree granting institutions. Only 42% of all faculty were women, 31.6% of tenured faculty were women and 43.7% of non-tenured faculty on the tenure track were women (2). Increasing the representation of women in science is firstly a matter of social justice, but secondly will potentially increase innovation and the impact of science. Mathematical modelling demonstrated that diverse groups can outperform homogeneous groups at problem solving (3). Experiments in problem solving by groups revealed a collective group intelligence (*c*)(analogous to an IQ for an individual) and one factor associated with *c* was the proportion of women in the group (4). In many fields, including business, racial and gender diversity increase success, e.g. in profits, customers and market share (5). In science, gender and/or ethnic diversity among authors increased the quality of journal of publication and the impact based upon citations of the paper and measures of novelty of the study (6-10). Balancing these benefits of increasing diversity are challenges resulting from the diversity of teams, but these can be effectively managed (11, 12).

To redress the issue of gender balance in the biomedical sciences it is critical to understand the underlying factors precluding equity in opportunity for men and women. Many studies have identified multiple factors that differentially impact men and women during their career trajectories, demonstrating complex causes and challenging solutions to overcome the imbalance in genders in the STEM fields. After completion of their PhD, men and women have different career interests (13). Surveys of postdocs and faculty who have left academia demonstrate that family factors are more important to women than to men (14, 15). Once established in an academic setting, retention between men and women is now relatively comparable (16), although the time until promotion is longer for women (17). Women receive fewer resources than men, including lower startup packages (18). While men and women generally have comparable success rates at securing extramural funding (19), men more frequently apply to renew grants, are more successful at renewing extramural grants and are more likely to have more funded projects at a time (20, 21). There is a difference in research output produced by men and women called the “productivity paradox”, which is discussed below. These and other factors differentially impact the pursuit of a scientific career by men and women.

There are established differences in publications, citations and patents between men and women (22-26). These differences are observed across disciplines and nationalities (23). Studies range from small studies focused upon a single institution, to entire fields of research (22, 27, 28), to large scale studies querying millions of publications (29, 30). These analyses are complicated by differences in career trajectories of men and women. One recent study documented a difference in total productivity but linked the difference in productivity to average length of publishing activity (31). On an annualized basis over periods of active publishing, the productivity of men and women was the same (31). This study underscores the importance of defining gender differences in performance at different career stages since the effect upon career outcome may be different, as can the factors impacting research productivity, at different career stages. Few publications have addressed research productivity of men and women doctoral students and these studies have focused upon single institutions (32-34). Analysis of the productivity of doctoral students is important since differences in research productivity at the very beginning of a scientific career can have a significant effect upon the trajectory of a scientific career (35). This study was undertaken to expand upon existing studies of doctoral student productivity by measuring research productivity of a large cohort of male and female doctoral students in the biological and biomedical sciences across many institutions in the US. The cohort contains more than 42,000 doctoral students at 235 institutions and finds a disparity in publications and citations between men and women. The findings underscore the necessity of addressing the “productivity paradox” amongst doctoral students as one strategy to solve the problem of disparity of women in the biomedical workforce.

## Methods

### Acquisition of Data

Data on a cohort of doctoral students in the biological and biomedical sciences completing their dissertations between 2012 and 2016 inclusive was collected. The names of doctoral students and advisors were collected from the ProQuest Dissertations & Theses Global database (access provided by West Virginia University libraries). Metadata from dissertations in the biological and biomedical sciences, as defined in the Graduate Students and Postdoctorates in Science and Engineering Survey (GSS), were retrieved. The data collected was restricted to doctoral students at doctorate granting institutions as defined in the GSS. Publication data was retrieved with the BioEntrez package from Python using the name of the doctoral student, the advisor and the university affiliation (36). To infer gender, a list of male names and female names was compiled from US Social Security Administration (SSA) data (30). The US SSA has compiled lists of male and female names of babies born each year that occur 5 or more times in the US (and the numbers of individuals with those names) since 1880 (https://www.ssa.gov/oact/babynames/limits.html). A complete list of names from 1880 through 2021 was assembled. Some names appear in both the list of boy’s names and girl’s names. Names where ≥ 90% of all individuals with that name belong to one gender were assigned to that gender. Names where < 90% of all individuals with that name belong to one gender were not assigned to a gender.

### Publications and Citations

First author publications were identified by searching PubMed for the doctoral student as first author, advisor as any author and the university affiliation. The metadata from ProQuest listed multiple advisors for many students. Individual PubMed searches using each listed advisor were performed and the results compiled. The search parameters for total publications were the doctoral student and advisor as any author and the university affiliation. The year of publication for each paper was captured and the number of citations for first author publications was determined using the BioEntrez package for Python (36). The PMID for each first author publication was used to search PubMed for the list of PMCIDs citing the publication and the number of PMCIDs tallied. Doctoral students were identified as male or female by matching their given name with the compiled lists of male names and female names.

### Institutional Review Board

The West Virginia University IRB approved the study. IRB approval number is WVU Protocol#: 2203537777.

### Statistics

Since none of the publication data exhibit a Gaussian distribution, nonparametric statistics were used for the analysis. Comparison of publications and citations between men and women were analyzed using the Mann Whitney test. Percentages of men and women publishing were compared using Fisher’s Exact test.

## Results

### Validation of the Names Database

The strategy to infer gender from given names utilized a names database collected from US Social Security Administration (SSA) data, similar to the approach used by West et al. (30) and was validated by the methods used by Santamaria and Mihaljevic, and Wais (37, 38). A successful gender inference strategy should correctly identify gender 98% of the time and have less than 25% unclassified (either not in the database or gender cannot be inferred, e.g. a name commonly used for males and females) (38). The names database assembled here was used to infer gender of names in a control dataset assembled from bibliographic records retrieved from zbMATH, genderizeR, PubMed and the Web of Science (39). The control dataset contains 7076 names with the gender independently verified for 5779 names. The names database assembled from SSA data contains unique male names, unique female names and names that are used for males and females. An arbitrary cutoff of 90% was used to infer gender to names that are used for males and females, i.e. if ≥ 90% of individuals with a name were of one gender, the name was used to infer that gender. If a name shared by males and females was used < 90% by one gender, that name was not assigned to a gender. The results of the validation analysis revealed that this strategy misidentified gender in 2.9% of the records. The percentage of unassigned names due to their absence from the names database was 18.2% and the percentage of unassigned names due to common use for males and females was 9.3%. There is a slight female gender bias (errorGenderBias = 0.019) (38). Adjustment of the cutoff value used to infer gender for common names for males and females changes these values. Increasing the stringency of the cutoff reduces misidentification, but increases the percentage of unassigned names. The 90% cutoff value was used to infer gender for the following analyses.

### Doctorates and Publications

Doctoral students completing dissertations in biological and biomedical sciences between 2012 and 2016 and their advisors were identified from the ProQuest Dissertations & Theses Global Database. Publications by the doctoral students were found by searching PubMed. The initial search provided publication records for 42,922 doctoral students who published 74,093 first author publications and 182,563 publications. This search strategy will include false positives due to more than one individual with the same name or publications by the doctoral student at a different stage in their education, e.g. as an undergraduate or postdoc. To partially correct for false positives the publications were restricted to a specific time frame and the top 1% of doctorates based upon number of publications were excluded as outliers. The differential between the year of publication of a paper and the year of the dissertations was determined (40). The 5^th^ percentile of first author publications was -3, i.e. 3 years before the dissertation year and the 95^th^ percentile was +4. For total publications, the 5^th^ percentile was -3 and the 95^th^ percentile was +6. The lower boundary for inclusion of publications was set at -3 (5^th^ percentile) and the upper boundary was arbitrarily set at +2 for first author publications and +3 for total publications. The rationale is to provide a reasonable time for completion and publications of studies following completion of the dissertation. The first author publication and total publication data were independently constrained and thus the two datasets do not consist of identical doctoral students. These restraints reduced the number of first authors to 42,481 doctoral students, who published 62,323 first author papers and 42,359 doctoral students with 128,813 total publications. Of the first author publications, 5,100 were reviews and 53,502 were primary peer-reviewed publications.

### Inferring Gender to Doctorates

The gender of the doctoral students was inferred from their given names, based upon the lists of male names and female names captured from the US SSA. In the doctoral students with first author publications 16,460 were inferred to be men and 16,457 were inferred to be women. A further 2,995 doctoral students had given names that could not be unambiguously assigned to a gender (7.1%). The remaining 6,568 doctoral students had given names that were neither in the list of boy’s names nor the list of girl’s names (15.5%). Thus, 84.5% of these doctoral students had given names captured from the US SSA data and gender was inferred for 77.5% of the doctorates. Fifty percent of the doctoral students were inferred to be women. Data from the Survey of Earned doctorates for 2012 to 2016 indicates that 53% of the students earning doctorates in the biological and biomedical sciences were women (1). Thus, the strategy applied to infer gender undercounts women. To ensure that the division of doctoral students into men and women did not alter the representation across institutions, the percentages of men and women doctoral students at different types of institution were compared with the entire cohort and were found to be similar (Supplemental Table 1).

A higher percentage of men published a first author paper than women (Table 1). The average number of first author papers published by men was 1.44 +/-1.38 compared with an average of 1.29 +/-1.30 papers published by women, a difference of 10.4%. Obviously, the fact that fewer women publish a first author paper contributed to the difference. Excluding all doctoral students who did not publish a first author paper, men published an average of 2.08 +/-1.18 papers and women published 1.98 +/-1.12 papers, a difference of 4.8%. A comparable percentage of men and women published first author reviews (∼ 11.2%).

**Table 1.** Research Productivity of Doctoral Students.

| **First Author Publications** |  |  |  |
| --- | --- | --- | --- |
|  | **Men** | **Women** | **p value** |
| % Published 1<sup>st</sup> Author | 69.1% | 65.2% | <0.0001 (F) |
| 1<sup>st</sup> Author Pubs | 1.44 +/- 1.38 | 1.29 +/- 1.30 | <0.0001 (MW) |
| 95% CI | 1.42 to 1.46 | 1.27 to 1.31 |  |
| Median 1<sup>st</sup> Author Pubs | 1 | 1 |  |
| 1<sup>st</sup> Author (Excl'd Zero) | 2.08 +/- 1.18 | 1.98 +/- 1.12 | <0.0001 (MW) |
| 95% CI | 2.06 to 2.10 | 1.95 to 2.00 |  |
| Median 1<sup>st</sup> Author Pubs (Excl'd Zero) | 2 | 2 |  |
| % Published 1<sup>st</sup> Author Review | 11.2% | 11.1% | 0.6115 (F) |
| **Citations** |  |  |  |
|  | **Men** | **Women** | **p value** |
| Avg Citations | 25.04 +/- 131.43 | 20.56 +/- 38.36 | <0.0001 (MW) |
| 95% CI | 23.36 to 26.71 | 20.04 to 21.07 |  |
| Median Citations | 12 | 11 |  |
| Avg Citations (Reviews) | 37.65 +/- 76.30 | 34.85 +/- 51.39 | 0.7315 |
| 95% CI | 34.36 to 40.94 | 32.60 to 37.11 |  |
| Median Citations (Reviews) | 20.5 | 21 |  |
| Avg Citations (Excl'd Reviews) | 23.83 +/- 135.5 | 19.07 +/- 36.43 | <0.0001 (MW) |
| 95% CI | 22.02 to 25.64 | 18.55 to 19.58 |  |
| Median Citations (Excl'd Reviews) | 11 | 11 |  |
| **Total Publications** |  |  |  |
|  | **Men** | **Women** | **p value** |
| % Published (total) | 79.8% | 76.4% | <0.0001 (F) |
| Total Pubs | 3.25 +/- 3.03 | 2.74 +/- 2.71 | <0.0001 (MW) |
| 95% CI | 3.20 to 3.29 | 2.70 to 2.78 |  |
| Median Total Pubs | 3 | 2 |  |
| Total Pubs (Excl'd Zero) | 4.07 +/- 2.85 | 3.59 +/- 2.57 | <0.0001 (MW) |
| 95% CI | 4.02 to 4.11 | 3.54 to 3.63 |  |
| Median Total Pubs (Excl'd Zero) | 3 | 3 |  |
| Time to 1<sup>st</sup> Pub | -1.24 +/- 1.47 | -1.02 +/- 1.51 | <0.0001 |
| 95% CI | -1.26 to -1.21 | -1.05 to -0.99 |  |
| Median Time to 1<sup>st</sup> Pub | -1 | -1 |  |

First author publications between years -3 and +2 (relative to year of dissertation), total publications between years -3 and +3 (relative to year of dissertation) and citations of first author publications. p value = the p value of a Mann Whitney test (MW) or a Fisher's Exact test (F).

In the analysis of total publications, 16,440 doctoral students were inferred to be men and 16,462 were inferred to be women (50%). The number of students who could not be unambiguously assigned to a gender was 2963. The remaining 6,494 students had given names that were not in the lists of male and female names. Approximately 77.7% of doctoral students could be assigned to a gender. The percentage of men and women from different types of institutions were comparable to the percentage of doctoral students distributed among different types of institutions in the entire cohort (Supplemental Table 2).

The percentage of men who published, including first author and co-author papers, was higher than the percentage of women who published (Table 1). Men published an average of 3.25 +/-3.03 total papers, while women published an average of 2.74 +/-2.71 papers. This is a difference of 15.8%. Excluding doctoral students, who did not publish, the average number of papers published by men was 4.07 +/-2.85 papers and the average number of papers published by women was 3.59 +/-2.57, a difference of 11.9%.

Men and women were also compared using two other metrics, citations and time until first publication. Citations are used as a surrogate for impact of the publication and time until first publication, relative to the dissertation year, is intended to compare the initial trajectory of careers. The average number of citations of men’s first author papers was 25.04 citations and the average number of citations of women’s first author papers was 20.56 (Table 1). On average, men’s first author papers receive 17.9% more citations than women’s first author papers. This difference was not due to different numbers of first author reviews written by men and women as the percentage of doctoral students publishing first author reviews was the same for men and women. Further, counting the citations for each first author review demonstrated equal numbers of citations for men and women (Table 1). The average time until first publication for men is - 1.24 years relative to the dissertation compare with -1.02 year for women’s first author publication. This is a difference of 2.64 months.

### Identification and Inferring Gender of Advisors

The meta-data associated with dissertations in the ProQuest database contains advisors for each doctoral student, but the names include only initials and surname. The author list of the publications of doctoral students were used to retrieve their advisors’ names, based upon the assumption that the senior author of a first author publication by a doctoral student is the student’s advisor. In the simplest scenario, there is one advisor linked to the dissertation and the advisor is a senior author on a first author publication by the doctoral student. This senior author is presumed to be the advisor for the analysis. If the advisor is not a senior author on the doctoral student’s first author publications, the advisor is considered ambiguous and the student is not matched with an advisor for the analysis. There are many instances where multiple advisors are listed in the dissertation. In the instances where a single advisor is a senior author on a first author publication by the student, this senior author is presumed to be the advisor for the analysis. In instances where the there are multiple advisors listed on the dissertation and more than one advisor is senior author on first author publications by the student, the assignment of an advisor is ambiguous and the student is not matched to an advisor for the analysis. A list of advisors who are matched to doctoral students at each institution was created. For doctoral students who have not published and have a single advisor affiliated with the dissertation and the advisor’s name matches a name on the list of advisors at that institution, the matched advisor is presumed to be the student’s advisor for the analysis. Advisor gender is inferred using the database of names created from US SSA data as above.

The number of unique advisors identified using this strategy was 16,670. Of these advisors, 10,107 were men and 3,796 were women (27.3% of individuals who were gender matched). There were 2,037 advisors’ whose given names were not in the database (12.3%) and 730 whose names could not be inferred using the gender inferring strategy (4.4%). According to the Survey of Doctoral Recipients from 2013 and 2015, the faculty (Professors/Associate Professors/Assistant Professors) in the broad discipline of Biological, Agricultural and Environment Life Sciences at four-year educational institutions consisted of 34% women. While this provides only a rough estimate of women faculty in the biological and biomedical sciences, it does suggest that women may be underrepresented in the sample in this analysis. Excluding advisors whose names could not be gender inferred, 73.7% of doctoral students worked with male advisors and 26.3% of the students had women advisors.

The 16,670 advisors were matched with 31,178 doctoral students. Of these students, the gender of 79.3% was inferred; 12,415 students were male and 12,312 students were female. The distribution of male and female doctoral students between male and female advisors differed. Excluding cases where either the doctorate or advisor could not be assigned a gender, male doctorates had male advisors in 77.9% of the cases and female advisors in 22.1% of the matches. Female doctorates had male advisors in 69.2% of the cases and female advisors 30.8% of the cases. The doctorates performing dissertations with male advisors were 52.8% male and 47.2% female, whereas doctorates performing dissertations with female advisors were 41.9% male and 58.2% female. These findings demonstrate a predilection for female students to work with female advisors.

The number of papers published by male and female doctoral students was recalculated from this dataset. Men published 1.82 +/-1.30 first author papers and women published 1.66 +/-1.26 (Table 2). The increased numbers relative to the data in Table 1 reflects a loss in doctorates without first author publications in the process of matching doctorates to advisors (e.g. 69.1% of men publish a first author paper in Table 1, whereas 86.3 of men publish a first author paper in the dataset of matched doctorates with advisors). Men’s first author papers were cited an average of 25.27 +/-134.0 times and women’s first author papers were cited 21.05 +/-47.16 times. In this dataset, male doctorates published 3.89 +/-2.89 total papers and female doctorates published 3.35 +/-2.62 papers. These results demonstrate similar differences in research productivity between men and women in this subset of doctoral students and the whole cohort.

**Table 2.** Research Productivity of Doctoral Students (Doctorate/Advisor Matched Dataset)

| **Doctorate Publications Based on Student Gender (Doctorate/Advisor Matched Dataset)** |  |  |  |
| --- | --- | --- | --- |
|  | **Men** | **Women** | **p value** |
| Number of Doctorates | 12,415 | 12,312 |  |
| 1<sup>st</sup> Author Pubs | 1.82 +/- 1.30 | 1.66 +/- 1.26 | <0.0001 (MW) |
| 95% CI | 1.80 to 1.85 | 1.64 to 1.69 |  |
| Median 1<sup>st</sup> Author Pubs | 2 | 1 |  |
| Number of Papers | 22,650 | 20,481 |  |
| Citations | 25.27 +/- 134.0 | 21.05 +/- 47.16 | <0.0001 (MW) |
| 95% CI | 23.52 to 27.01 | 20.40 to 21.69 |  |
| Median Citations | 12 | 11 |  |
| Number of Doctorates | 12,267 | 12,242 |  |
| Total Pubs | 3.89 +/- 2.89 | 3.35 +/- 2.62 | <0.0001 (MW) |
| 95% CI | 3.84 to 3.94 | 3.30 to 3.39 |  |
| Median Total Pubs | 3 | 3 |  |
| **Doctorate Publications Based on Gender of the Advisor (Matched Dataset)** |  |  |  |
|  | **Men** | **Women** | **p value** |
| Number of Advisors | 19,476 | 6,942 |  |
| 1<sup>st</sup> Author Pubs | 1.74 +/- 1.30 | 1.78 +/- 1.29 | = 0.0023 |
| 95% CI | 1.72 to 1.76 | 1.75 to 1.82 |  |
| Median 1<sup>st</sup> Author Pubs | 2 | 2 |  |
| Number of Papers | 33,820 | 12,387 |  |
| Citations | 24.43 +/- 117.40 | 20.45 +/- 33.20 | = 0.1494 |
| 95% CI | 23.18 to 25.68 | 19.86 to 21.03 |  |
| Median Citations | 12 | 12 |  |
| Number of Advisors | 19,289 | 6,898 |  |
| Total Pubs | 3.64 +/- 2.80 | 3.52 +/- 2.60 | = 0.1228 |
| 95% CI | 3.60 to 3.68 | 3.46 to 3.58 |  |
| Median Total Pubs | 3 | 3 |  |

First author publications between years -3 and +2 (relative to year of dissertation), total publications between years -3 and +3 (relative to year of dissertation) and citations of first author publications. p value = the p value of a Mann Whitney test (MW) or a Fisher's Exact test (F).

The impact of the gender of the advisor upon the publication record of their doctoral students was of interest. The average number of first author publications by doctoral students (regardless of gender) performing dissertations with male advisors was 1.74 +/-1.30 (Table 2). The average numbers of first author publications by doctoral students working with female advisors was 1.78 +/-1.29. The number of citations of first author papers from male advisors was 24.43 +/-117.4 and the number of citations of first author papers from female advisors was 20.45 +/-33.20. The number of total publications by doctoral students with male advisors was 3.64 +/-2.80 and for students with female advisors was 3.52 +/-2.60. The difference in the number of first author publications was statistically different and students working with women advisors had more publications. The differences in citations and in total publications was not significantly different. Given the literature describing the difference in research productivity between men and women, it is particularly intriguing that doctoral students working with male advisors exhibit research productivity equal to doctoral students working with female advisors.

In the cases where gender could be inferred for both the doctoral students and advisors, student:advisor dyads were analyzed for research productivity (Table 3). Female doctoral students working with male advisors published the fewest average number of first author publications (1.63 +/-1.24) compared with the other three dyads (1.73 +/-1.3 to 1.81 +/-1.24). The difference was statistically significant. Examining the students working with female advisors exclusively showed that male students published significantly more first author papers than female students. This difference paralleled the observations in the entire male and female cohorts. The number of first author papers was not significantly different between all male dyads and all female dyads. Dyads with male doctoral students published more total papers (3.72 +/-2.64 and 3.90 +/-2.91) than dyads with female doctoral students (3.33 +/-2.63 and 3.33 +/-2.53) regardless of the gender of the advisor. Again, these results are completely in accord with the results of the entire cohort of male and female doctoral students. The female:female dyads had significantly fewer citations per first author paper (19.12 +/-30.73) than any other dyad, while the male:male dyads had the most citations per first author paper (27.02 +/-161.2) than the others. This observation infers that male:male dyads produce papers with greater impact, while female:female dyads produce papers with the least impact.

**Table 3.** Research productivity of student:advisor dyads.

|  | **Dyads** |  |  |  |
| --- | --- | --- | --- | --- |
|  | **MM** | **MF** | **FM** | **FF** |
| Number | 8256 | 2358 | 7381 | 3277 |
| 1<sup>st</sup> Author Pubs | 1.79 +/- 1.31 | 1.81 +/- 1.24 | 1.63 +/-1.24** | 1.73 +/- 1.30* |
| 95% CI | 1.76 to 1.82 | 1.76 to 1.86 | 1.60 to 1.66 | 1.69 to 1.78 |
| Median 1<sup>st</sup> Author Pubs | 2 | 2 | 1 | 2 |
| KW = 76.23, p<0.0001; **MM>FM (p<0.0001), MF>FM (p<0.0001), FF>FM (p=0.0018), *MF>FF (p=0.0158) |  |  |  |  |
|  | **MM** | **MF** | **FM** | **FF** |
| Total Pubs | 3.90 +/- 2.91 | 3.72 +/- 2.64 | 3.33 +/- 2.63* | 3.33 +/- 2.53* |
| 95% CI | 3.83 to 3.96 | 3.62 to 3.83 | 3.27 to 3.39 | 3.24 to 3.41 |
| Median Total Pubs | 3 | 3 | 3 | 3 |
| KW = 196.1, p < 0.0001; *MM = MF > FM = FF (p<0.0001) |  |  |  |  |
|  | **MM** | **MF** | **FM** | **FF** |
| Avg Citations | 27.02 +/- 161.2 | 22.63 +/- 36.08 | 22.22 +/- 54.25 | 19.12 +/- 30.73 |
| 95% CI | 24.45 to 29.58 | 21.55 to 23.70 | 21.26 to 23.19 | 18.32 to 19.91 |
| Median Citations | 12 | 13 | 12 | 11 |
| KW = 39.02, p<0.0001; MM>MF (p=0.0444), MM=FM, MM>FF(p<0.0001), MF>FM (p=0.0009), MF > FF (p<0.0001), FM>FF (p=0.0072) |  |  |  |  |

First author publications between years -3 and +2 (relative to year of dissertation), total publications between years -3 and +3 (relative to year of dissertation) and citations of first author publications. p value = the p value of a Mann Whitney test (MW) or a Fisher's Exact test (F).

## Discussion

The results of this study demonstrate a gender disparity in research productivity of doctoral students in a large national cohort of students. Men publish 10.4% more first author papers and 15.7% more total papers than women (Table 1). First author papers by male doctoral students receive 17.9% more citations that first author papers written by female doctorate students (Table 1). Several other studies have examined differences in research productivity between male and female doctoral students, although the studies were restricted to single elite universities (32, 34). Both studies found a similar disparity between the research productivity of men and women. The first of these studies examined a cohort of 933 doctoral students in Engineering and Natural Sciences and found that women doctoral students published 8.5% fewer papers than their male counterparts (32). The second was a survey-based study of 1285 recent doctoral graduates from a wide range of fields (34). The men in this cohort published an average of 4.9 papers and the women published an average of 2.9 papers, a publication difference of 40%. A third study of a small national cohort of first year graduate students in the biological sciences demonstrated that for every 100 hours worked, men were 15% more likely to publish than women (33). While the magnitude of the difference differs, the conclusion is consistent across these studies. The current study establishes the generality of these observations in a large national cohort containing a wide range of institutions.

Interestingly, the research productivity of all doctoral students working for male advisors is comparable to the productivity of all doctoral students working for female advisors. In fact, students with female advisors publish slightly more first author papers. This result is surprising, given the established disparity in gender productivity between men and women (22). As the current study focused on a narrow window of time, the observation is consistent with the view that annualization of research productivity eliminates differences in research productivity of men and women over their careers (31). Pezzoni et al. also examined the gender of advisors for their cohort (32). They captured 204 unique advisors, 12.3% of whom were women. Women were advisors for 11% of the students. Overall, male advisors published more papers per year than women advisors (7.6 papers vs 6.3 papers). To contrast, the current study identified 16,670 unique advisors, 83.4% of whom had a gender inferred. Of these advisors, 27.3% were women and these were advisors for 26.3% of the doctoral students. Both studies concur that a higher percentage of female doctoral students work with advisors who are women than male doctoral students. Both studies also agree that the research productivity of doctoral students working with female advisors is equal to or exceeded the research productivity of doctoral students working with male advisors. However, Pezzoni et al. found that the overall research productivity of male advisors exceeded that of female advisors (32). Thus, the disparity in total publications between male and female advisors disappeared if the analysis is restricted to the publications of their doctoral students.

Pezzoni et al. report that male doctoral students working with female advisors publish 10% more papers than male students working with male advisors, who in turn publish 8.5% more papers than female students working with male advisors (32). The productivity of male:male and female:female dyads were similar. The results in Table 3 show a similar trend when comparing first author publications, but this trend is not observed when comparing total publications. This result suggests that the dyad affects publications from projects led by the doctoral student, but not other projects where the student participates as a member of a team.

## Limitations

This study utilizes the ProQuest Dissertations & Theses Global Database to identify doctoral students and PubMed to identify publications. The appropriate and most rigorous method for gender identification is self-identification, which is not possible using these large public databases. Further, this study uses binary gender identification based upon given names, which is an oversimplification of the complexity of gender identity. A US centric database of names was utilized for gender matching, which is appropriate for the majority of the students and faculty at US institutions, but misses accurate gender matching for students and faculty whose given names are not frequently found in the US Social Security Administration database. The study relies upon the identification of publication records of doctoral students based upon names and affiliations. The dataset will contain false positives in cases where individuals with the same name have the same affiliation as a doctoral student identified from the ProQuest Dissertations & Theses Global Database. Despite efforts to reduce the number of false positives by constraining publications by time and excluding records with very high numbers of publications, false positives may persist. In addition to false positives, the dataset is also missing data. Complete dissertation information is not available for all institutions in the ProQuest Dissertations & Theses Global Database. These limitations in the dataset are not anticipated to impact the outcomes of the study.

### Potential Consequences of Productivity Disparity

Differences in research productivity as a doctoral student could affect career development over both the short and long term. Differences in productivity as graduate students could be one explanation of why there are fewer women who train as postdoctoral fellows in elite labs (41). Publication record, even at the very beginning of a research career, is predictive for becoming a principal investigator (35), and could be a factor contributing to the disparity of men and women in the biomedical workforce. Individuals with early publications are obviously more competitive candidates for future positions, but early research success also has broader effects. Either the factors producing the differences in research productivity or early productivity itself could impact the research self-efficacy of junior scientists. Research self-efficacy and first author publication rate correlates with interest in pursuing an academic career (13) and higher first author publication rates correlate negatively with interest in other careers. Thus, the gap in research productivity could have direct and indirect effects upon the successful pursuit of a career in science. Interventions to close this gap or policies to accommodate this gap may be necessary to ensure equal opportunities for men and women to advance beyond the entry level into a scientific career.

### Potential Causes of Productivity Disparity

A number of survey-based studies have attempted to address the underlying causes resulting in publishing disparity between men and women in graduate school. The disparities in publishing parallel disparities in submissions of manuscripts (34, 42). Women doctoral students in their first year report more time working on research than their male counterparts (33). The tasks required to gain authorship may be different for men and women. In attributing contributions to papers, women are more likely to report performing experiments than men, whereas men are more likely to report designing experiments, analyzing data and contributing to manuscript writing than women (43). Importantly, these differences are observed even at the beginning of publishing careers, i.e. at the time of doctoral studies (43). First year male doctoral students report greater confidence in their abilities to develop hypotheses and design experiments (33). In survey responses, men express a higher satisfaction with their relationship with their mentor and in preparation for their career and these factors are significant positive predictors of submissions (34). In one study, men more frequently report that their mentor encourages them to publish than women and this is a positive predictor of submission (34). In contrast, a broad survey across programs at a single university reports little difference in response by men and women to a prompt asking if their mentor encouraged publication (42). Finally, women are more likely to report that their progress toward publication was impaired by family obligations and faculty availability and these factors were negative predictors of manuscript submissions (34). The potential underlying causes of the disparity in research productivity between men and women are varied as will be strategies to address the disparity. Further, disparities can differ even between programs at the same institution (42), demonstrating differences in underlying factors even at the local level. Based upon the circumstances and climate at different institutions and in different programs, strategies to address gender disparities in productivity will need to be tailored to the specific circumstances.

