## Supplemental Material for "The Gender Gap Amongst Doctoral Students in the Biomedical Sciences"

**Supplemental Table 1. Distribution of Doctoral Students (1st Authors) at Different Types of Institution**

| | Cohort | Men | Women |
|---|---|---|---|
| At Private Unis | 34.3% | 34.2% | 34.8% |
| At Public Unis | 65.7% | 65.8% | 65.2% |
| At Land Grant Unis | 30.0% | 29.9% | 29.3% |
| At IDeA Unis | 10.4% | 10.2% | 10.5% |
| At Carnegie Highest | 80.6% | 81.0% | 80.7% |
| At Carnegie Higher | 12.9% | 12.8% | 12.5% |
| At Carnegie Special | 4.3% | 4.3% | 4.8% |

Public, private and land grant institutions were identified according to the Graduate Students and Postdoctorates in Science and Engineering Survey (GSS). Institutions in IDeA states were identified using the NIH definition. Carnegie status in 2015, i.e. within the time frame of analysis, is according to the GSS and uses the Carnegie definitions at that time.

**Supplemental Table 2. Distribution of Doctoral Students (Total Authors) at Different Types of Institutions**

| | Cohort | Men | Women |
|---|---|---|---|
| At Private Unis | 34.3% | 34.1% | 34.8% |
| At Public Unis | 65.7% | 65.9% | 65.2% |
| At Land Grant Unis | 30.1% | 29.9% | 29.3% |
| At IDeA Unis | 10.4% | 10.3% | 10.5% |
| At Carnegie Highest | 80.6% | 81.0% | 80.6% |
| At Carnegie Higher | 13.0% | 12.8% | 12.6% |
| At Carnegie Special | 4.3% | 4.3% | 4.8% |

Public, private and land grant institutions were identified according to the Graduate Students and Postdoctorates in Science and Engineering Survey (GSS). Institutions in IDeA states were identified using the NIH definition. Carnegie status in 2015, i.e. within the time frame of analysis, is according to the GSS and uses the Carnegie definitions at that time.
